# Vulnerability of invasive glioma cells to lysosomal membrane instabilization

**DOI:** 10.1101/276402

**Authors:** Vadim Le Joncour, Maija Hyvönen, Pauliina Filppu, Pauliina S. Turunen, Harri Sihto, Isabel Burghardt, Heikki Joensuu, Olli Tynninen, Juha Jääskeläinen, Michael Weller, Kaisa Lehti, Pirjo Laakkonen

**Affiliations:** Research Programs Unit, Translational Cancer Biology, University of Helsinki, Helsinki, 00290, Finland; Research Programs Unit, Genome-Scale Biology, University of Helsinki, Helsinki, 00290, Finland; Department of Microbiology, Tumor and Cell Biology (MTC) Karolinska Institutet, SE-17177 Stockholm, Sweden; Department of Neurology and Brain Tumor Center, University Hospital Zurich and University of Zurich, Zurich, 8091, Switzerland; Department of Oncology, Helsinki University Hospital, Helsinki, 00029, Finland; Department of Pathology, Haartman Institute, University of Helsinki and HUSLAB, Helsinki, 00014, Finland; Kuopio University Hospital, Kuopio, 70210, Finland; Laboratory Animal Centre, University of Helsinki, Finland

**Keywords:** glioma, LMP, anti-histamine, cell death, MDGI

## Abstract

Diffusive by nature, glioma challenges clinical care by the impossibility of complete surgical resection of tumor, leaving the radio- and chemoresistant cells responsible for recurrence intact. We identified mammary-derived growth inhibitor (MDGI/*FABP3*) as invasive glioma biomarker. Here, we show that high MDGI expression associated with poor patient survival and promoted invasive glioma cell growth both *in vitro* and *in vivo*, while MDGI silencing drastically compromised patient-derived tumoroid viability via induction of lysosomal membrane permeabilization (LMP). This alternative cell death pathway provokes release of lysosomal hydrolases into the cytosol leading inevitably to the cell death. Our results show a novel functional role for MDGI in glioma cell invasion, survival, and maintenance of the lysosomal membrane integrity as well as an unsuspected sensitivity of glioma cells to an LMP-inducing drug, anti-histamine clemastine. In a preclinical study, clemastine-treatment significantly prolonged the survival of intracranial glioblastoma-bearing animals due to eradication of invasive glioma cells. This glioma cell vulnerability to LMP-inducing drugs opens new horizons for development of novel treatments and suggest re-positioning of an established drug for new indication.

## Introduction

Gliomas constitute approximately 30% of all primary nervous system tumors and 80% of all malignant brain tumors (1). Glioblastoma is the most frequent, aggressive, and lethal type of gliomas. Glioblastomas harbor a dense abnormal vasculature, display large hypoxic and necrotic areas, and contain extensively proliferating tumor cells with the intrinsic ability to disseminate and colonize the brain far beyond the primary tumor mass. The current standard of care, comprising surgery, radio- and chemotherapy, provides only modest improvement in the patient survival, and the prognosis remains dismal (2). This is due to *(i)* impossibility of complete surgical resection of the tumor; *(ii)* intratumoral heterogeneity and presence of multidrug-resistant cells and stem cell-like glioma cells responsible for the tumor maintenance and relapse, and *(iii)* the sheltering effect of the blood-brain-barrier (BBB), which efficiently prevents administration of many systemic anti-cancer agents into the brain. Most probably different approaches are required to eradicate the invasive cells and the cells that reside within the tumor bulk (3). Thus, novel therapeutic approaches for glioblastomas are urgently needed (4).

We have previously identified mammary-derived growth inhibitor (MDGI) as a glioma biomarker expressed in tumor cells and their associated vasculature (5). MDGI, also known as heart-type fatty acid binding protein (H-FABP/*FABP3*), belongs to the family of fatty acid binding proteins (FABPs) that facilitate the intracellular transport of fatty acids (6). Both tumor-suppressive (7) and tumor-promoting (8, 9) functions, depending on the cancer type, have been reported for MDGI. In glioma cells, MDGI has been found to mediate lipid droplet formation and palmitate uptake (10).

Here, we used cohorts of patients operated for a primary glioma and novel patient-derived human tumoroid cultures to examine the function of MDGI. MDGI was abundantly expressed in clinical gliomas, where its expression was associated with poor survival of patients. MDGI was upregulated by hypoxia and its overexpression enhanced the invasive growth of glioma cells both *in vitro* and *in vivo*. Surprisingly, MDGI silencing compromised tumoroid growth and glioma cell survival via lysosomal membrane permeabilization (LMP). Accordingly, we show that glioma cells were more sensitive than normal cells to an LMP inducing drug, the anti-histamine clemastine, *in vitro* and that clemastine treatment was able to eradicate the invasive glioma cells *in vivo*. Our results suggest that MDGI expression is crucial for glioma cell viability and an important regulator of lysosomal integrity. These characteristics make MDGI a potential therapeutic target and LMP-inducing drugs a promising treatment option to eradicate especially the invasive glioma cells.

## Results

### High MDGI expression correlates with poor survival

Our previous results show that MDGI is highly expressed in glioblastomas (5). To study the potential correlation of MDGI expression with the clinicopathological variables and patient survival, we performed immunohistochemistry for MDGI in human tumor microarrays (TMAs) consisting of lower WHO grade (grade II-III) gliomas and glioblastomas and scored the staining intensity separately in tumor cells and tumor-associated endothelial cells. Approximately 50% of both grade II-III gliomas and glioblastomas expressed moderate to high levels of MDGI, accompanied with positive vascular staining for MDGI (Appendix Tables S1-2). Only 5% of all gliomas (n = 6/122) lacked detectable MDGI expression. In the glioblastoma specimens, MDGI expression correlated with the presence of the CD117/C-Kit-receptor in the perinecrotic tumor regions (*P*=0.006) (Appendix Table S2). No association was detected between MDGI expression and EGFR (p>0.999), EGFRvIII (*P=*0.613), phosphorylated EGFR (Tyr-1173) (p>0.999) or p53 (*P*=0.499) (Appendix Table S2). Among the lower grade gliomas, MDGI expression did not correlate with the sex, tumor grade or the WHO 2007 histological type (p > 0.5; Appendix Table S1).

Moderate to high MDGI levels significantly associated both with poor glioma (grade II-III) -specific (HR = 1.85; 95% CI: 1.08-3.16; *P=*0.022; Appendix Figure S1A) and poor overall survival of patients (HR = 1.98; 95% CI: 1.19-3.28; *P=*0.007, Figure 1A) as compared to patients with negative or low tumor MDGI expression. Multivariable Cox hazards analysis showed that both MDGI expression and high tumor grade were independently associated with unfavorable overall survival, increasing the risk of death by the factor of 2 (Table 1). In glioblastomas, tumor MDGI levels were not associated with overall survival (Appendix Figure S1B). In addition to the patient tissue biopsies, MDGI expression was high in seven distinct patient-derived tumoroid cultures, whereas it was low in all five adherent cell lines studied (Figure 1B).

**Table 1.** Cox multivariant analysis of association of MDGI expression and tumor grade on glioma patient survival.

| Covariate | $\beta$ (SE) | HR | *P* |
| --- | --- | --- | --- |
| MDGI expression<br>Positive vs. negative | 0.56 (0.264) | 1.76 (1.05 to 2.94) | 0.033 |
| Tumor grade<br>III vs. II | 0.76 (0.264) | 2.14 (1.28 to 3.60) | 0.004 |

**Figure 1.**
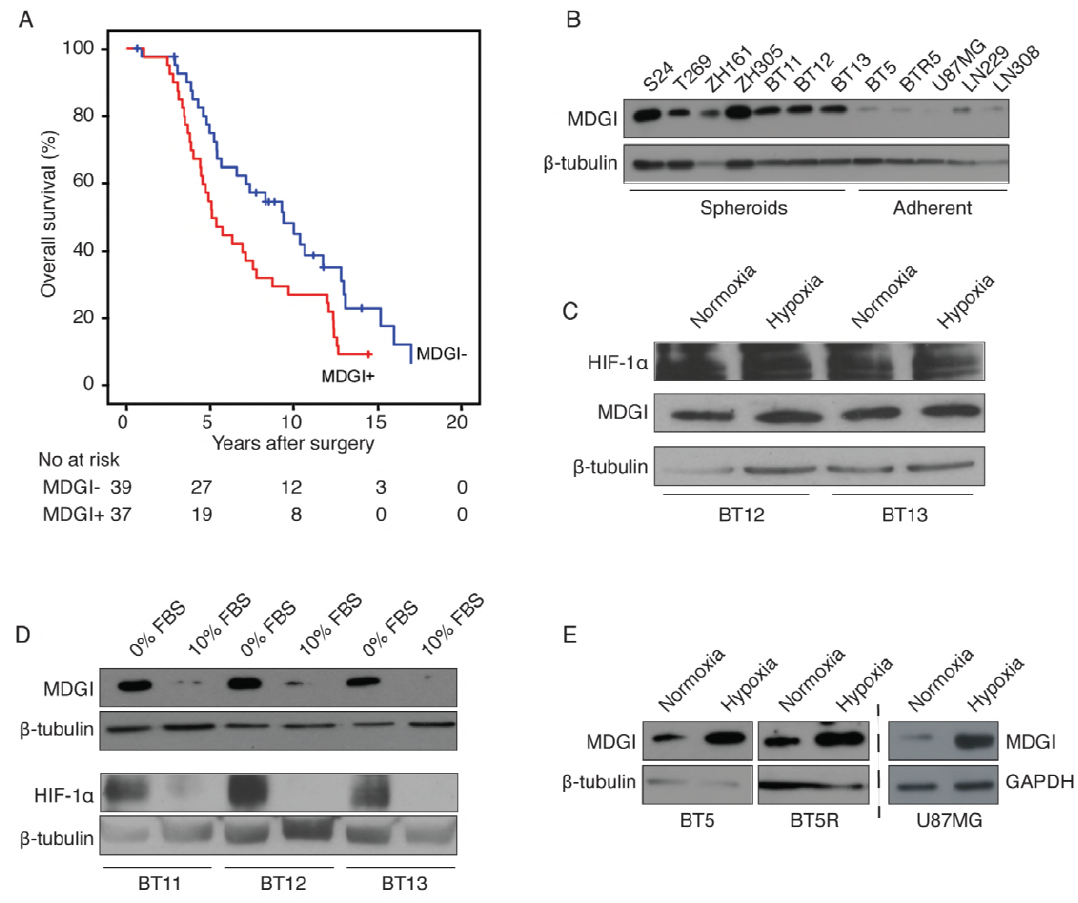
Association of MDGI protein expression with patient outcome and regulation of MDGI expression by hypoxia. A, Overall survival profile of grade II and III glioma patients (n=76) was significantly better in patients with none/low (blue line) compared to patients with moderate/high (red line) MDGI expression (*P*=0.007). The cumulative survival rates of patients were estimated by using the Kaplan– Meier method. B, Western blot analysis shows MDGI expression in human glioma cells including patient-derived tumoroids (S-24, T269, ZH161, ZH305, BT11, BT12, BT13), adherent, patient-derived (BT5 and BT5R), and long-term cell lines (U87MG, LN229, LN308). C, Expression of MDGI and HIF-1α in tumoroids after 24h culture under hypoxia (1% O_2_) or normoxia. D, Expression of MDGI and HIF-1α in tumoroids after seven days of culture in medium containing none or 10% of serum (FBS). E, MDGI expression in adherent glioma cell lines after 24h culture under hypoxia or normoxia. In B-E representative Western blot images of 2-3 separate experiments are shown and β-tubulin served as loading control.

### MDGI overexpression promotes glioma cell invasion

Our immunohistochemical results in clinical tumor samples revealed a correlation between MDGI expression and perinecrotic C-Kit, which is an indirect hypoxia marker in glioblastomas (11). In patient-derived BT12 and BT13 tumoroids, MDGI expression was high in conjunction with strong expression of hypoxia-inducible factor 1 α (HIF-1α) regardless of the culture condition under normoxia or hypoxia (Figure 1C). However, serum-containing medium, which shifted the HIF-1α expressing hypoxic tumoroids to adherent monolayer growth, reduced both MDGI and HIF-1α levels (Figure 1D). Moreover, MDGI expression was highly increased by hypoxia in adherent cell lines (Figure 1E), linking MDGI induction in glioma cells closely to hypoxia.

As MDGI expression was associated with poor prognosis in the glioma patient cohort, we studied its function in glioma cell growth and invasion. We overexpressed MDGI as a GFP-fusion protein (MDGI-GFP) in the U87MG cells since they express low endogenous levels of MDGI and form local, non-invasive tumors following intracranial injection in pre-clinical models. While MDGI overexpression did not affect cell proliferation (Appendix Figure S1C), it significantly enhanced colony formation, suggesting an increased capacity for aggressive, anchorage-independent growth of the MDGI overexpressing cells (Figure 2A-B). In addition, MDGI overexpressing tumoroids grew more invasively in an *ex vivo* brain slice model compared to the control cells (Figure 2C-D). Moreover, the intracranial U87MG-MDGI-GFP xenografts grew invasively (Figure 2E-F), formed satellite tumors in the brain (Figure 2G-H, K), and displayed vascular co-option (Figure 2I-J, L) unlike the control GFP-expressing U87MG derived xenografts that, as expected, only formed cyst-like delineated masses.

**Figure 2.**
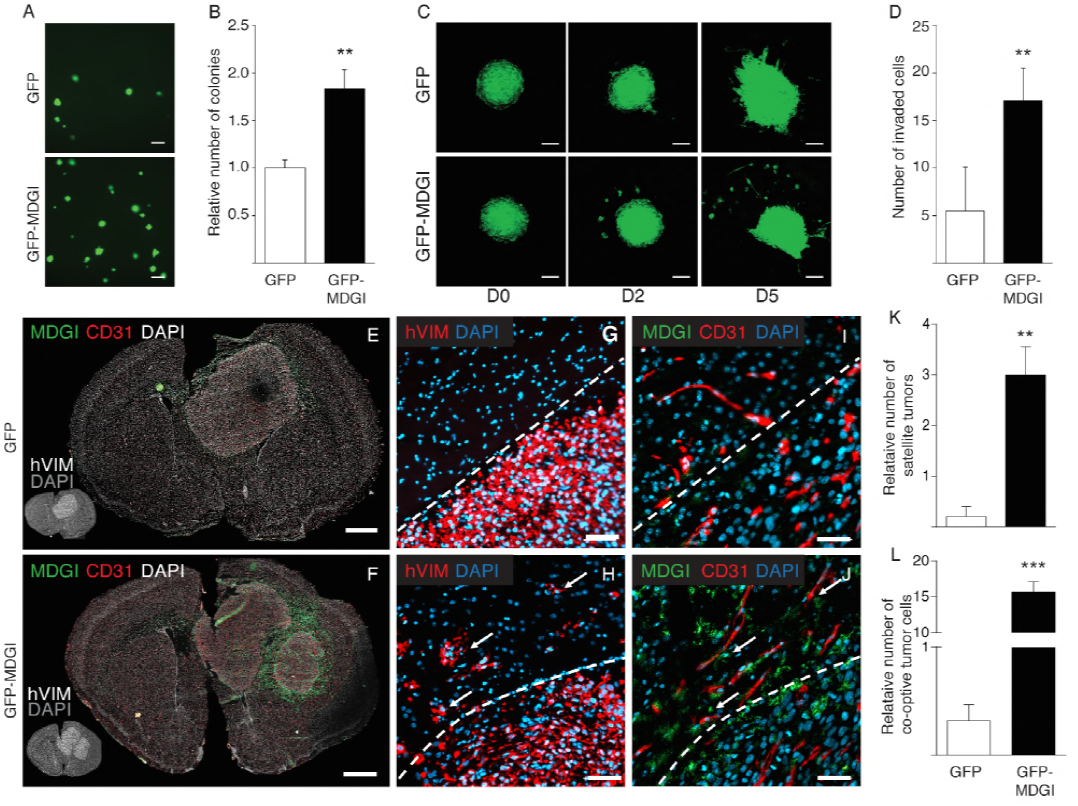
MDGI expression promotes the aggressive and invasive growth of glioma cells. A, Colony formation assay demonstrates the anchorage-independent growth of the U87MG cells expressing MDGI-GFP or GFP after 14 days of culture. Scale bar: 200 μm. B, Quantification of the number of GFP-MDGI colonies (solid bar) relative to the GFP control (open bar; n=3). C, Invasion/migration of U87MG glioma cells expressing MDGI-GFP or GFP on murine brain sections *ex vivo*. Spheres were placed on top of brain slices and their motility was recorded by confocal microscopy at the indicated time points (D0, D2 and D5, n≥7). Each microphotograph is the maximal intensity projection from 10 μm thick optical slices (18-22 slices per sample). Scale bar: 200 μm. D, Quantification of the number of invaded MDGI-GFP cells (solid bar) relative to the GFP controls (open bar; n≥7). E-F, Representative microphotographs of whole murine brain coronal sections (bregma +0.98 mm) injected with GFP (E) or GFP-MDGI (F) expressing U87MG glioma cells. Xenografted human cells were visualized by using anti-human vimentin (thumbnail, hVim white), MDGI using anti-MDGI (green) and blood vessels using anti-CD31 antibodies (red). Scale bar: 500 μm. G-J, Microphotographs of consecutive (separating distance: 9 μm) brain sections stained for human glioma cells using antibodies specific for human vimentin (hVim red in G and H), MDGI (green in I and J) and CD31 (red in I and J). Dashed lines separate the tumor and the healthy brain. Arrows in H indicate angiotropic, co-opting tumor cells (red) expressing MDGI (green in J) next to blood vessels (red in J). Scale bar: 50 μm. K, Quantification of the number of satellite tumors (diameter >300 μm) detected in the whole brain (GFP: n=5, GFP-MDGI: n=9). L, Quantification of the number of co-opting tumor cells detected in the brain (GFP: n=5, GFP-MDGI: n=9). Data are represented as mean ± SD in B and D and as mean ± SEM in K and L. **, *P*<0.01; ***, *P*<0.001. P-values were calculated using two-tailed, nonparametric Mann-Whitney *U* test.

### MDGI silencing dramatically reduces glioblastoma cell viability

To confirm MDGI’s importance in aggressive cell growth, we silenced MDGI in in patient-derived BT12 and BT13 glioblastoma cells using shRNA (shMDGI1). Efficient gene silencing was verified using Western blot analysis (Appendix Figure S1D). MDGI silencing caused a dramatic change in the cell morphology and disappearance of the large multicellular tumoroids (Figure 3A). Moreover, the anchorage-independent growth of the MDGI silenced cells was severely compromised (Figure 3B). Surprisingly, MDGI silencing inhibited proliferation of both BT12 and BT13 cells (Figure 3C) and dramatically reduced their viability (Figure 3D). Similar, but slightly less drastic effects were obtained with another shRNA-construct (shMDGI2) with approximately 70-80% knockdown efficiency (Appendix Figure S1E-G and S2A-B); demonstrating a dose-dependent effect of MDGI silencing on glioblastoma cell growth and viability. Next, we studied whether MDGI silencing induced cell death was affected by the EGFR expression, since *EGFR* is mutated, amplified, or both in about 60% of glioblastomas. Only the patient-derived glioblastoma BT11, BT12, and BT13 tumoroids expressed high levels of EGFR, while expression was low in U87MG and LN308 cells (Figure 3E). No EGFR expression was detected in other cell lines studied (Figure 3E). MDGI silencing induced death also of the ZH305 tumoroids that did not express EGFR indicating that EGFR status of glioma cells did not affect their sensitivity to LMP (Figure 3F-G).

**Figure 3.**
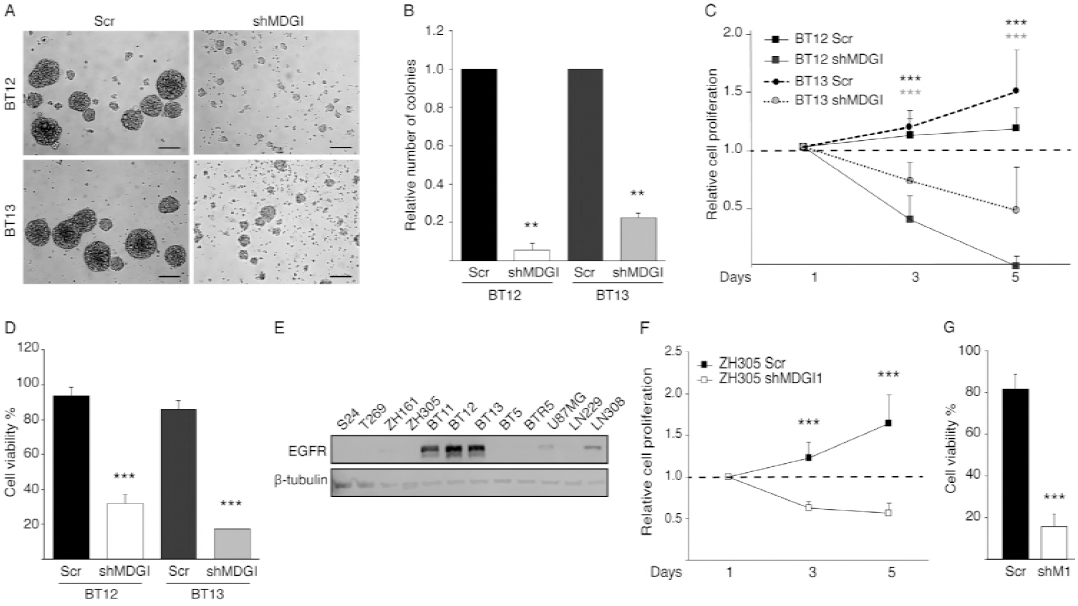
MDGI silencing decreases glioma cell growth and viability. A, Light microscopic images of MDGI-silenced (shMDGI) and control (Scr) patient-derived BT12 and BT13 tumoroids 11 days after gene silencing. Representative images of more than 3 independent experiments are shown. Scale bar: 200 μm. B, Colony formation assay demonstrating the anchorage-independent growth of control and MDGI-silenced patient-derived BT12 and BT13 cells after 27 days of culture. The number of colonies relative to the controls is indicated (n=3). C, MTT-proliferation assay of control and MDGI-silenced cells. The values of each cell line were normalized to the day 1 values (n=3). D, Cell viability was determined using the Trypan blue staining 10 days after MDGI silencing (n=3). E, Western blot analysis shows EGFR expression in human glioma cell lines. β-tubulin served as loading control. F, MTT-proliferation assay of control and MDGI silenced ZH305 cells. The values of each cell line were normalized to the day 1 values (n=3). G, Cell viability was determined using the Trypan blue staining 10 days after MDGI silencing (n=3). Data are represented as mean ± SD. **, *P*<0.01. ***, *P*<0.001. P-values were calculated using two-tailed, nonparametric Mann-Whitney U test.

### MDGI silencing induces caspase-independent glioma cell death

To further examine the mechanism underlying the reduced viability of MDGI silenced cells, we studied the binding of Annexin V to the exposed phosphatidylserines as a marker of cell death (12). We observed significantly increased binding of Annexin V to the MDGI silenced cells indicating increased cell death (Figure 4A-B and Appendix Figure S2C-D). To study the intracellular pathways that could contribute to this increased cell death, we analyzed the expression of apoptosis-associated proteins (Figure 4C and Appendix Figure S2E) at various time points after lentiviral MDGI silencing. Expression of the phosphorylated and total p53 as well as the pro-apoptotic protein BAD remained first stable and were eventually decreased 5-6 days after transduction (Figure 4C and Appendix Figure S2E). Surprisingly, the amounts of total caspase 3 and its cleaved, activated form were also decreased 5-6 days after silencing (Figure 4C and Appendix Figure S2E). This suggests that apoptosis in the MDGI silenced cells was not mediated by the caspase activation. To verify the results, we next performed an antibody array of apoptosis-associated proteins using extracts of control and MDGI silenced cells. When fold-changes less than 0.6 or more than 1.5 compared to control was used as cut-offs, levels of the anti-apoptotic survivin and X-linked inhibitor of apoptosis protein (XIAP) were decreased. On the other hand, the protein levels of the HTRA serine protease were elevated (Appendix Figure S3A-B). The levels of various caspases, pro-apoptotic proteins BAD and BAX or cytochrome C were not changed by MDGI silencing (Appendix Figure S3A-B). Furthermore, MDGI silencing-induced cell death could not be rescued by silencing of the pro-apoptotic protein BAD by siRNA (Appendix Figure S3C-D).

**Figure 4.**
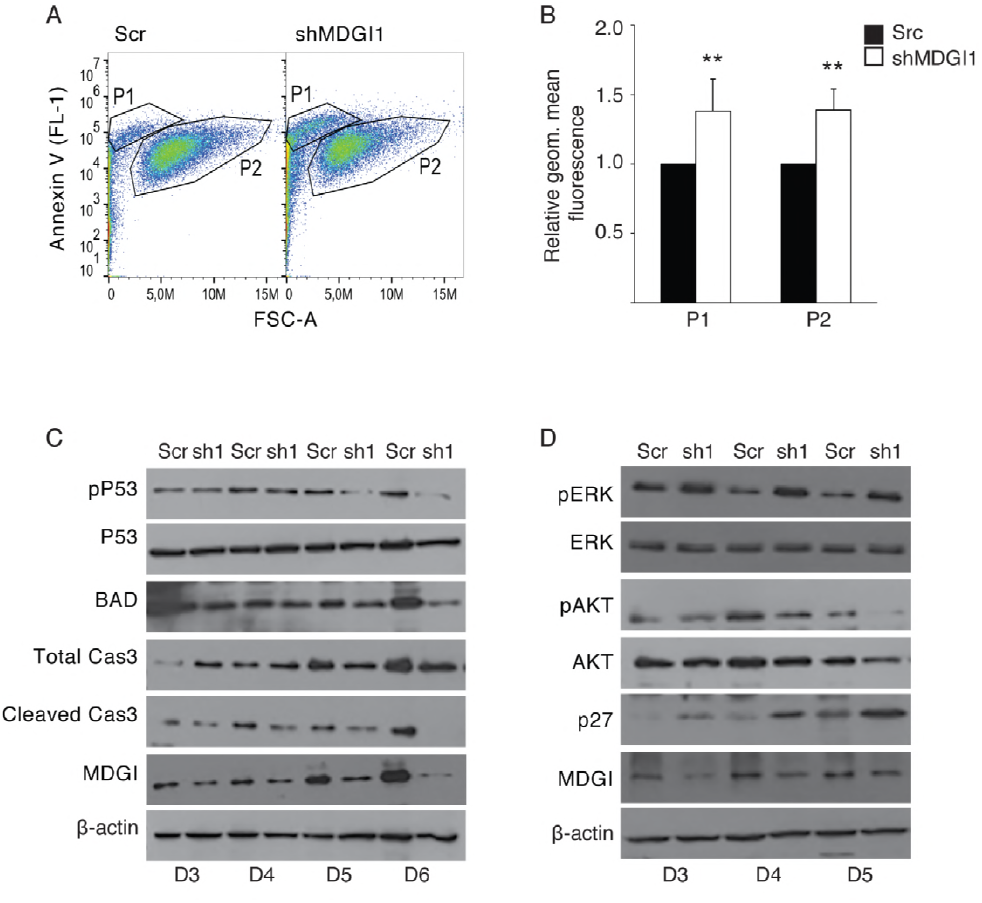
MDGI silencing induces cell death. A, Binding of Annexin V Alexa Fluor 488 to cells was measured by flow cytometric analyses five days after MDGI silencing (shMDGI1). The analysis was performed by gating the cells into two populations: P1 (small, granular) and P2 (normal-sized cells) (30 000 analyzed events/cell line, n=3). B, Graph shows the results as geometric mean fluorescence (FL-1) relative to the controls in P1 and P2 in MDGI silenced (shMDGI1) and control (Scr) cells. C, Western blot analysis shows expression of apoptosis-associated proteins after MDGI silencing. Cell extracts were collected at indicated time points from control (Scr) and MDGI silenced (sh1) cells. Representative images of 2 individual experiments are shown. D, Western blot analysis shows expression of selected intracellular signaling proteins after MDGI silencing. Cell extracts were collected at indicated time points from control (Scr) and MDGI silenced (sh1) cells. Representative images of 2 individual experiments are shown. Data are represented as mean ± SD. **, *P*<0.01. P-values were calculated using two-tailed, nonparametric Mann-Whitney U test.

We also studied the expression of selected proteins of the key signaling pathways at different time points after MDGI silencing using Western blot (Figure 4D and Appendix Figure S2F). In MDGI silenced cells, p27 levels were elevated and the Erk1/2 phosphorylation was significantly induced, while the phosphorylated AKT was decreased (Figure 4E and Appendix Figure S2F) already at day three after silencing.

### MDGI silencing induces lysosomal membrane permeabilization (LMP)

Since apoptosis in MDGI silenced cells seemed not to be mediated by caspase activation, and their phenotype could not be rescued by silencing of the pro-apoptotic protein BAD, we studied the effects of MDGI silencing on LMP. This alternative cell death pathway leads to the release of lysosomal hydrolases into the cytosol and can, depending on the extent of the release, ultimately lead to lysosomal cell death with necrotic or apoptotic features (13). LMP induction can be easily visualized as a change in galectin-1 localization from a diffuse cytoplasmic to a punctate staining pattern (14). As a control, we first treated the tumoroids with the LMP-inducing agent, L-leucine O-methyl (LLOMe) (15), and detected increased formation of galectin-1 (LGALS1) -positive puncta (Figure 5A-B). Similarly, we detected a significant increase in the number of LGALS1-positive punctate staining in the MDGI silenced patient-derived BT12 and BT13 cells (Figure 5C-G). In addition, a significant fraction of the MDGI silenced cells was already dead at the time of analysis, as judged by small and fragmented nuclei and no detectable LGALS1-staining (Figure 5H). LGALS1-puncta co-localized with the lysosomal marker protein, lysosome-associated membrane protein 2 (LAMP2), verifying the lysosomal localization of LGALS1 (Figure 5I-K and Appendix Figure S4A-C). Moreover, the activity of lysosomal cathepsin B was increased in the cytoplasm of the MDGI silenced cells (Figure 5L) confirming the destabilization of the lysosomal membranes.

**Figure 5.**
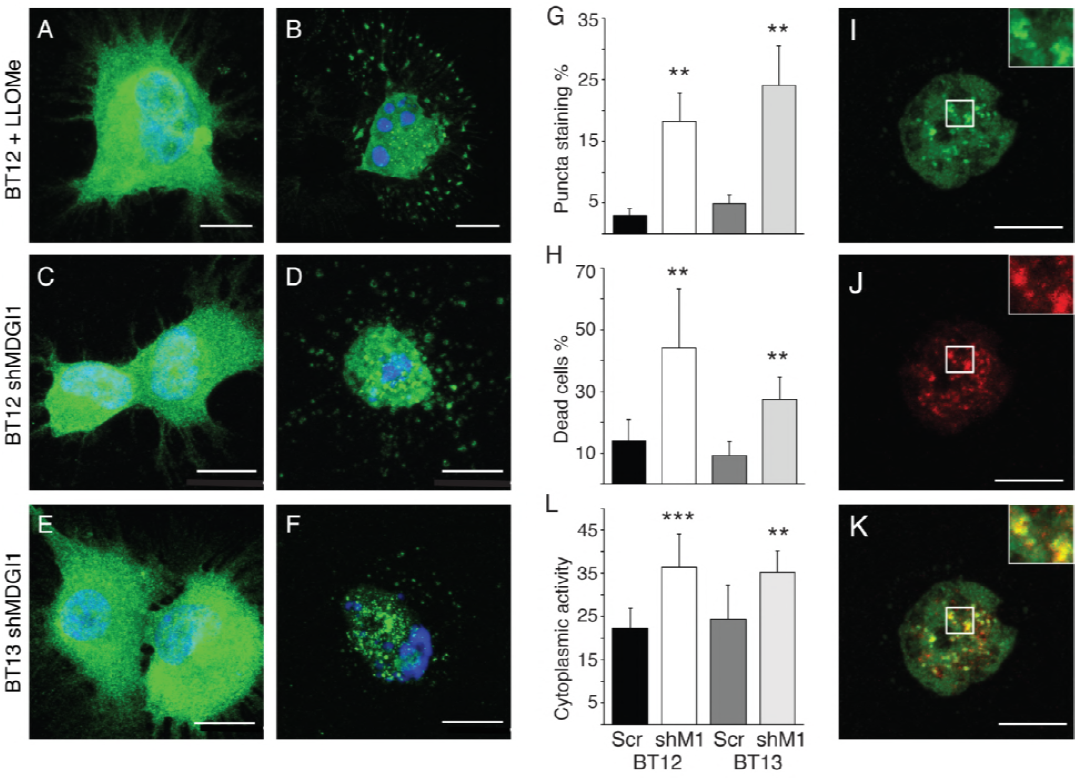
MDGI silencing induces glioma cell death via lysosomal membrane permeabilization (LMP). A-B, Representative images of BT12 non-treated control cells (A) and cells treated with 2 mM LLOMe for 4 hrs (B) before staining by using the anti-galectin-1 (LGALS1) antibody (green). C-D, Representative images of LGALS1-stained (green) BT12 control (C) and MDGI silenced cells (D) six days after silencing. E-F, Representative images of LGALS1-stained (green) BT13 control (E) and MDGI silenced cells (F) six days after silencing. Nuclei were visualized by using DAPI (blue) in A-F. G, Graph demonstrates the percentage of LGALS1-positive puncta staining in the control (Scr) and MDGI silenced (shM1) cells. In total 0.4-1.2×10^4^ LGALS1-stained cells were analyzed from 50 mm^2^ coverslip areas (n=6). H, Quantification of the percentage of unstained, dead cells six days after MDGI silencing and LGALS1 staining (n=100) in control (Scr) and MDGI silenced (shM1) cells. I-K, Co-localization of the LGALS1 (green in I) with the lysosomal marker protein LAMP2 (red in J) is seen as yellow color (K) in MDGI silenced BT12 cells. Scale bar: 10 μm. Data are represented as mean ± SD. L, Graph shows the cytoplasmic cathepsin B activity in the control (Scr) and MDGI silenced (shM1) cells. **, *P*<0.01; ***, *P*<0.001. P-values were calculated using two-tailed, nonparametric Mann-Whitney U test.

### Clemastine evokes glioma cell death

Inspired by MDGI silencing-induced cell death via LMP, we searched for an LMP-triggering drug that could be safely used in a pre-clinical study. A recent report by Ellegaard et al. identified cationic amphiphilic (CAD) anti-histamines as drugs able to induce LMP (16). Therefore, we chose clemastine (Tavegil™), a first-generation histamine H1 blocking antihistamine CAD, as the BBB-permeable drug for our experiments. The patient-derived BT12 and BT13 glioblastoma cells were treated with increasing concentrations of clemastine (1-5 μM). Cell viability was decreased by 90% and 50% after 96 hrs of treatment at 2 μM and 1 μM clemastine concentrations, respectively, (Figure 6A-B). No significant cell death was observed when normal human (HuAR2T) or murine (bEND3) endothelial or embryonic kidney (HEK293T) cells were treated at the same concentrations (Figure 6C-D, Appendix Figure S4D) suggesting a therapeutic window for clemastine treatment in gliomas. In accordance, already 1 μM of clemastine induced punctate localization of the galectin-1 in BT12 and BT13 cells (Figure 6E) whereas no re-localization of galectin-1 was observed in HuAR2T and HEK293T cells (Figure 6F). Galectin-1 relocation into the lysosomes in BT12 and BT13 cells was confirmed by co-localization with the LAMP2 (Figure 6G-I).

**Figure 6.**
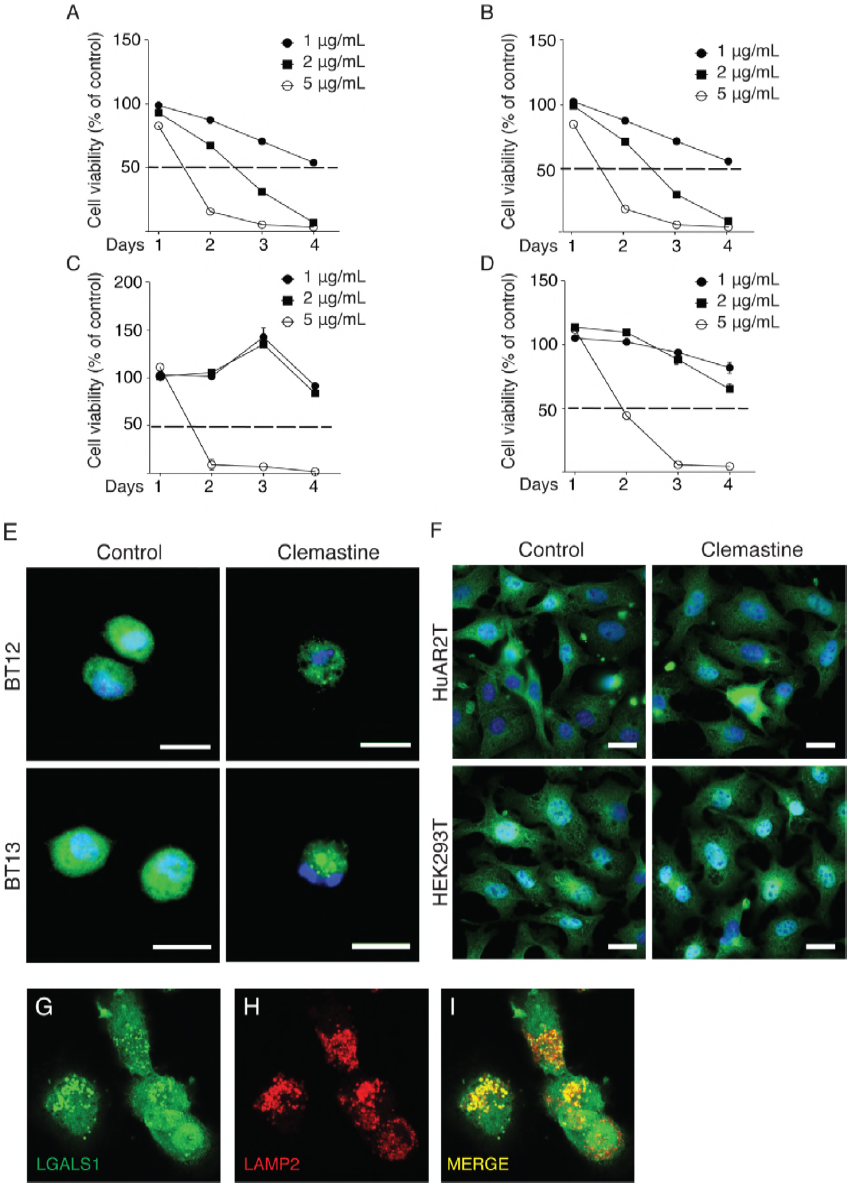
Anti-histamine treatment induces glioma cell death via lysosomal membrane permeabilization (LMP). A-B, Measurement of the BT12 (A) and BT13 (B) glioblastoma cell viability at the indicated clemastine concentrations and time points (n=12). C-D, Measurement of the HuAR2T (C) and HEK293T (D) cell viability at the indicated clemastine concentrations and time points (n=12). A dashed line in A-D marks the 50% cell viability. E, Representative images of BT12 and BT13 glioblastoma cells treated with 1 μg of clemastine for 48h and stained with the anti-galectin-1 (LGALS1) antibody (green). Non-treated cells served as control. Nuclei were visualized by using DAPI (blue). Scale bar: 20 μm. F, Representative images of normal human endothelial (HuAR2T) and kidney (HEK293T) cells treated with 1 μg clemastine for 48h and stained with the anti-LGALS1 antibody (green). Non-treated cells served as control. Nuclei were visualized by using DAPI (blue). Scale bar: 10 μm. G-I, Co-localization of the LGALS1 (green in G) with the lysosomal marker protein LAMP2 (red in H) is seen as yellow color (I) in clemastine treated BT12 cells. Data are represented as mean ± SD. **, *P*<0.01; ***, *P*<0.001. P-values were calculated using two-tailed, nonparametric Mann-Whitney U test.

The preclinical evaluation of clemastine was next performed in patient-derived xenografts orthotopically implanted in immuno-compromised mice. After 15 days of tumor growth, we started intraperitoneal injections of clemastine at a dose of 100 mg/kg on the first day followed by 50 mg/kg daily injections for 12 additional days. The saline solution used as a vehicle was administered under the same modalities to the control cohort. As illustrated in the Figure 7A, clemastine provoked a profound alteration in the xenograft growth pattern. The number of invasive tumor satellites was dramatically reduced (Figure 7B). As a consequence, clemastine treated animals showed significantly prolonged survival (*P=*0.044) compared to the controls (Figure 7C). In addition, clemastine treatment significantly reduced the number of invasive cells that had escaped the primary tumor and disseminated into the brain (Figure 7D, F-G). To further evaluate the antitumor effect of clemastine we analyzed the number of apoptotic cells by terminal deoxynucleotidyl transferase dUTP nick end labeling (TUNEL) in control and clemastine treated animals. Whole brain quantification revealed that the leading, migratory edge of the tumor was more susceptible to the cytotoxic effect of clemastine (Figure 7H-J) than the tumor cells inside the tumor mass. In addition, clemastine-treatment led to a nearly complete loss of co-opting tumor cells (Figure 7E and K).

**Figure 7.**
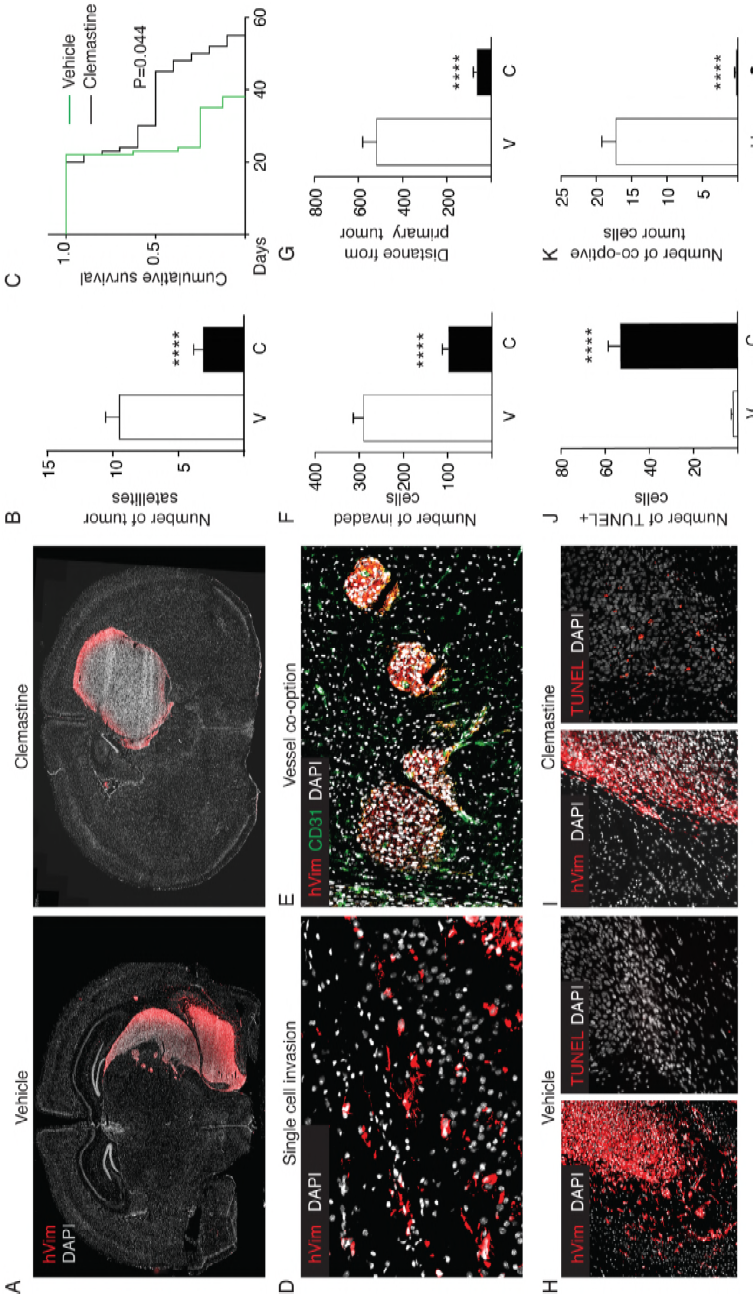
Clemastine treatment of intracranial patient-derived glioblastomas inhibits tumor cell invasion and co-option of blood vessels. A, Representative microphotographs of whole mouse brain coronal sections (Vehicle bregma −1.94, clemastine −1.06; mm) intracranially implanted with BT12 glioblastoma cells and daily treated with saline vehicle or 50 mg/kg clemastine for 12 days. Tumor cells were visualized by staining with an anti-vimentin antibody specific for human vimentin (hVim, red). Scale bar: 500 μm. B, Quantification of the number of the invasive tumor satellites in the brains of vehicle (open bars) and clemastine treated (solid bars) animals (Vehicle n=9, clemastine n=12). C, Kaplan-Meier survival curve of the BT12 xenografted animals treated with vehicle (green line, n=15) and clemastine (black line, n=12). D, Representative microphotographs of the invaded single glioblastoma cells (red) along the mouse *corpus callosum*. Nuclei were visualized by using DAPI (white). E, Representative microphotograph of the angiotropic glioblastoma cells (red) co-opting murine brain microcapillaries (green). F, Quantification of the number of the invaded single cells in the murine brain for the indicated treatments (V=Vehicle: n=9, C=clemastine: n=12). G, Quantification of the distance of the single invaded glioblastoma cells from the primary tumor (V=Vehicle: n=9, C=clemastine: n=12). H, Representative microphotographs of consecutive brain sections (separating distance: 9 μm) xenografts treated with saline vehicle and labeled with the anti-human vimentin (red in left panel) and TUNEL (red in right panel). Scale bar: 50 μm. I, Representative microphotographs of consecutive brain sections (separating distance: 9 μm) of xenografts treated with clemastine and labeled with the anti-human vimentin (left, red) and TUNEL (middle, red). Scale bar: 50 μm. J, Quantification of the number of TUNEL positive cells in the murine brain for the indicated treatments (V=Vehicle: n=9, C=clemastine: n=12). K, Quantification of the number of angiotropic tumor cells that have co-opted blood vessels in the murine brain for the indicated treatments (V=Vehicle: n=9, C=clemastine: n=12). Data are represented as mean ± SEM. ****, *P*<0.0001. P-values were calculated using two-tailed, nonparametric Mann-Whitney *U* test.

## Discussion

There is an unmet need for novel therapeutic strategies to treat gliomas making the characterization of proteins involved in disease progression highly important. Here we examined the expression of a potential glioma biomarker, MDGI (5), in clinical grade II-III glioma and glioblastoma specimens as well as in patient-derived and commercially available glioma cell lines. We show that high MDGI expression associated with unfavorable patient survival and was crucial for glioma cell viability. We further describe a novel function for MDGI in the maintenance of lysosomal membrane integrity and provide data that shows the exceptional vulnerability of invasive glioma cells to destabilization of lysosomes.

MDGI was frequently expressed in human gliomas and showed a significant correlation with poor survival. High MDGI expression was also detected in various patient-derived, cancer stem cell-enriched tumoroids but not in adherent cells under normoxia. However, hypoxia could stimulate MDGI expression in adherent cells. This is in agreement with previous data that showed upregulation of MDGI expression during hypoxia (10). MDGI expression in adherent glioma cells significantly promoted their anchorage-independent growth and invasion both *in vitro* and *in vivo*, suggesting a functional role for MDGI in the invasive growth. Although MDGI expression has been linked to tumor suppressive properties in breast cancer (7), in gastric carcinomas MDGI associated with poor patient survival (8) and in melanomas its expression was upregulated during disease progression (9) suggesting that the effect of MDGI on tumorigenesis may be tissue and cancer type-dependent.

Interestingly, our results demonstrate that MDGI expression is crucial for glioma cell survival. One of the central genes promoting glioblastoma cell survival is EGFR (17). It addition, breast cancer cell resistance to EGFR targeted therapy has been shown to be mediated by MDGI (18). However, it appears that the cell death induced by MDGI silencing is independent of the EGFR status of glioma cells. The ZH305 cells that do not express EGFR show the similar drastic loss of cell viability after MDGI silencing when compared to the BT12 and BT13 tumoroids expressing high amounts of EGFR. In addition, we observed significantly increased LMP in the MDGI silenced cells. LMP is a cell death pathway that leads inevitably to cell death due to irreversible leakage of lysosomal enzymes to the cytoplasm, where they digest vital proteins and intracellular organelles (13). Due to MDGI’s fatty acid binding and transport ability, it is possible that its silencing may lead to alterations in the lysosomal membrane composition. Previous report showed that incorporation of arachidonic acid and choline glycerophospholipid into phospholipid membranes is reduced in the MDGI knockout mice (19). In addition, activation of Erk in the MDGI silenced cells could play a role in the process, since phosphorylated Erk -besides being linked to apoptosis-induction and cell cycle arrest (20)-has also been shown to enhance cysteine cathepsin expression and activity, sensitizing cells to LMP (21). In addition to the LMP, we also observed decreased activity of the AKT kinase (the master regulator of various cell survival pathways (22)), downregulation of anti-apoptotic proteins XIAP and survivin, which has been linked to caspase-independent apoptosis (23) as well as an upregulation of the pro-apoptotic serine protease HTRA (24) and cyclin-dependent kinase inhibitor p27/Kip1, a negative regulator of the cell cycle (25). HTRA has been shown to contribute to cell death by downregulating XIAP protein levels (26) and decrease chemoresistance of colon cancer cells (27).

LMP induction has been shown effective against apoptosis-resistant cancer cells and re-sensitize multidrug resistant cells to chemotherapy (28, 29). LMP induction in glioblastoma treatment has not been widely studied, but several studies have recently characterized the mechanism of action of some novel chemotherapeutics and shown that lysosomal dysfunction plays a major role in drug-mediated cancer cell death (30–32). Among the drugs currently on the market, cationic amphiphilic (CAD) anti-histamines induce lysosomal cell death. Moreover, use of the CAD antihistamines was associated with significantly reduced all-cause mortality among cancer patients when compared with the use of non-CAD antihistamines and adjusted for potential confounders (16). We chose an older generation CAD anti-histamine, clemastine, due to its ability to cross the BBB, to test the sensitivity of our patient-derived glioblastoma cells towards LMP induction. Clemastine is devoid of neurotoxic effects (33) even though it causes reported fatigue-induced side effects common to most of the anti-histamine drugs (34). It has been shown to exert regenerative properties on the optic nerve (35), induce re-myelinization (36) and temper down inflammation (37) in amyotrophic lateral/multiple sclerosis preclinical models. Moreover, clemastine is FDA-approved and currently under clinical evaluation for the above mentioned neurodegenerative disease (38). We observed a dramatic loss of the glioblastoma cell viability that was associated with the loss of lysosomal membrane integrity at doses that did not affect the proliferation or viability of several normal cells *in vitro*. Surprisingly, when we evaluated the pre-clinical efficacy of clemastine, the survival of animals bearing intracranial glioblastoma xenografts was significantly prolonged compared to controls due to the eradication of invasive glioma cells. Both co-optive single cell invasion and collective migration of glioblastoma cells were inhibited by the systemic delivery of clemastine leading to a nearly complete loss of disseminated cells that form the clinical challenge in glioma treatment. It has become clear that different treatment modalities are required for the cells within the tumor bulk and the invasive cells. The cytotoxic effect of clemastine targeted preferentially the invasive cells. As most of the systematically administered chemotherapies (39), clemastine delivery into the primary tumor mass was likely challenged by the poorly functional tumor blood vessel network (40). The resistance of the tumor bulk towards clemastine may also be due to the chemoattraction of glioma cells to subventricular/intraventricular space (41) that was observed in some of our animals or the extreme resistance of glioma cells when part of a connective network compared to single cells (42).

Many triggers of the LMP are known (13, 43, 44), but genes and cellular pathways that regulate the lysosomal membrane integrity remain poorly understood. Our study demonstrates the crucial role of MDGI in glioma cell survival and its involvement in the LMP, suggesting a novel role for MDGI in maintaining lysosomal membrane integrity. A very recent study by Guishard et al. showed the existence of an evident translational gap in the past and ongoing clinical glioma trials due to the lack of attempts to include drugs that are able to cross the BBB and overcome the chemoresistance of glioma stem cells (3). The inoperable and highly invasive and chemo-resistant glioma stem cell-like cells showed an unexpected fragility towards LMP. Thus, providing a potential answer to the clinical challenge of the management of the cells that eventually will lead to the recurrence of the disease. These results also suggest re-positioning of an old drug with a new indication. More widely, LMP-inducing agents should be considered as a possible novel treatment option for gliomas.

## Materials and methods

### Glioma patient series

The glioblastoma and grade II-III glioma patient biopsies were obtained from the Department of Pathology, Helsinki University Hospital, Finland. The series have been described elsewhere (45, 46). Briefly, glioblastoma TMA was composed of formalin-fixed paraffin embedded (FFPE) tumors derived from 43 craniotomy patients, subsequently diagnosed with primary glioblastoma between October 2000 and December 2003. The median survival time of the patients was 9 months (range 1-40 months) as calculated from the date of the diagnosis to the date of death. All patients died due to glioblastoma during follow-up. In seven cases, tissue microarray sample was missing or staining was not informative leaving 36 patients in the series.

FFPE grade II-III glioma TMAs were prepared from the tumors of 122 patients who were operated for a primary glioma at the Helsinki University Hospital between 1979 to 2000. In ten cases, tissue microarray sample was missing or staining was not informative leaving 112 patients for the analysis. 33 of the tumors (30%) were diagnosed as astrocytoma, 16 (14%) as anaplastic astrocytoma, 36 (32%) as glioblastoma, 18 (16%) as oligodendroglioma and 9 (8%) as oligoastrocytoma. The association of MDGI with clinical and histopathological factors was studied from 76 low-grade samples, excluding glioblastoma patients and the median follow-up time after surgery was 6.0 years (range 0.6-17.0 years). Ten of the patients lived at the end of the follow-up, while 58 and 8 patients died from glioma or other causes, respectively.

### Immunohistochemistry and evaluation of immunostaining

The detailed protocol is found in Appendix Supplementary Material. MDGI expression was evaluated blinded to the clinical data and scored as follows: 0, all tumor cells negative; 1, low expression in tumor cells (5% to 50%); 2, moderate expression in ≥50% of tumor cells; and 3, high expression in ≥50% of tumor cells. For the statistical analyses, 0 and 1, as well as 2 and 3 tumors were grouped together. MDGI’s vascular expression was scored as follows: 0, no MDGI-positive vessels; 1, one or more MDGI-positive vessels.

### Patient-derived cells

The glioma patient samples were obtained from surgeries (Kuopio University Hospital, Finland) during years 2010-2011. BT5 cells were obtained from a human gliosarcoma and BT5R cells from its relapse. BT11 cells are from a glioblastoma-containing oligocomponents, BT12 and BT13 cells are from glioblastoma patients. Protocol for the cell isolation can be found in the Appendix Supplementary Material.

### Cell culture

The patient-derived brain tumor cells (BT11, BT12, BT13, BT5, BT5R) were maintained for up to 20 passages in Dulbecco’s Modified Eagle Medium with Nutrient Mixture F-12 (DMEM/F12, Gibco) and patient-derived cell line ZH305 (generated at Zurich University Hospital) was maintained for up to 10 passages in phenol red-free Neurobasal medium (Invitrogen). Both media were supplemented with 2 mM L-glutamine, 2% B27-supplement (Gibco), 100 U/ml penicillin and 100 μg/ml streptomycin, 0.01 μg/ml human fibroblast growth factor (FGF, Peprotech), 0.02 μ/ml human epidermal growth factor (EGF, Peprotech), and 15 mM HEPES-buffer. All the cells were maintained and grown at 37°C in a humidified atmosphere containing 5% CO_2_ unless stated otherwise. Mycoplasma contamination was routinely checked twice a month using the mycoplasma detection kit (11-1050, Minerva Labs). Additional information about the commercial cell lines can be found in the Appendix Supplementary Material.

### Generation of MDGI silenced cells

The MDGI/*FABP3* shRNA-constructs in the pLKO.1 vector were obtained from the RNAi Consortium shRNA library (Broad Institute of MIT and Harvard). The sequences were as follows: shMDGI1: ^3’^GACCAAGCCTACCACAATCAT^3’^, shMDGI2: ^5’^GACAGGAAGGTCAAGTCCATT^3’^. Lentivirus production and transduction protocol can be found in the Appendix Supplementary Material.

### Anchorage-independent growth assay

Cells (3000/35-mm well, triplicates) were suspended in medium containing 0.35% agarose and pipetted into 6-well plates containing a 2-ml layer of solidified 0.7% agar in the medium. Complete medium was added twice a week. After 14 (U87MG) or 27 (BT12 and BT13) days, eight images per well were taken (Zeiss Axiovert) and the number of colonies was quantified using the ImageJ software (National Institute of Health, USA). The experiment was repeated twice.

### *Ex vivo* brain slice cultures

Female FVB mice (5-8 weeks of age) were anesthetized and their brains were removed. Cerebellum was excised and the two hemispheres were embedded into 4% Low Melting Point Agarose (Thermo Scientific) in PBS. Slices of 500 μm were cut with a vibratome (HistoLab) and placed on top of 0.4 μm filter membranes (Millipore) in 6-well plates. Neurobasal A (1 ml, Gibco) supplemented with 1 mM glutamine, 2% B-27 (Gibco) and 1% penicillin/streptomycin was added to the bottom of each well. The medium was changed every other day. The slices were incubated for one week (37°C, 5% CO_2_) before starting the experiments (47). The spheres were formed o/n in U-shaped 96-well plates coated with 0.6% agarose (4000 cells/well), after which they were placed to the brain slices. The GFP-fluorescent spheres were imaged at indicated time points by taking 10 μm optical slices with Leica TCS SP2 confocal microscope. Image stacks were generated using the ImageJ-software and the number of invaded and migrated cells/cell groups were quantified manually. The experiment was repeated twice with 3-5 individual spheres/cell line/slice in each experiment.

### Analysis of lysosomal membrane permeabilization

LMP was determined using the LGALS1 puncta-staining assay (14). BT12 cells treated with 2 mM L-leucine O-methyl ester (LLOMe, Santa Cruz Biotechnology) were used as positive controls for LMP. For the quantification of LGALS1 staining (cytoplasmic/puncta), DAPI and LGALS1-stained cells grown on PDL-coated coverslips were scanned with 3DHISTECH Pannoramic P250 FLASH II using Zeiss Plan-Apochromat objective (20x/NA 0.8) and pico. Edge 4.2 CMOS camera. The number of LGALS1-stained cells (cytoplasmic/puncta) and unstained dead cells were manually calculated from each coverslip (n=6, 50 mm^2^ area) with Pannoramic viewer-software (3DHISTECH.)

### LMP induction by clemastine

Clemastine fumarate salt (Sigma) cytotoxicity was tested at the indicated concentrations and time points using the MTT protocol as described in the Appendix Supplementary Material. LMP induction by clemastine was verified using the LGALS1 puncta-staining assay described in the previous section (14).

### Animal experiments

Animal experiments were approved by the Committee for Animal Experiments of the District of Southern Finland (ESAVI/6285/04.10.07/2014). Six-weeks-old immunocompromised NMRI-nu (Rj:NMRI-Foxn1^nu^/Foxn1^nu^, Janvier Labs) mice were intracranially engrafted using a stereotaxic injector (World Precision Instruments) in the corpus callosum with 10^5^ U87MG-GFP (n=5) or U87MG-GFP-MDGI (n=9) cells in 5 μl of PBS under ketamine and xylazine anesthesia. Post-operative painkiller (temgesic) was locally administered for two days. Following the same methodology, the clemastine preclinical study included the implantation of 10^5^ BT12 cells dissociated from tumoroids in 5 μl of PBS. Two weeks after tumor implantation, animals were randomized and clemastine (200 μl in PBS, intraperitoneally, n=10) was injected daily at the indicated doses for a duration of 12 days in one cohort or until the physical manifestation of tumor burden for the survival studies. Control animals (n=15) were treated under the same modalities with the vehicle (200 μl). At the endpoint of the experiment, animals were euthanized and brains were snap-frozen in −50°C isopentane (Honeywell) until tissue processing as described in the Appendix Supplementary Material.

### Patient study approval

The ethics committee of the Hospital District of Helsinki and Uusimaa approved this study. The Ministry of Social Affairs and Health, Finland, permitted the use of tumor tissue. The ethics committee of the Pohjois-Savo Health Care District municipalities permitted the use of human glioma tissue specimens (53/2009, 1.8.2009-31.7.2019).

### Statistics

Association of MDGI expression with other factors was tested using a Chi-squared test or Fisher’s exact test and Mann-Whitney’s *U*-test was used for the MDGI vs age analysis. All tests were 2-tailed. The cumulative survival rates of patients were estimated by using the Kaplan–Meier method, and survival between groups, hazard ratio (HR), and their 95% confidence interval (95% CI) were computed by using the Cox proportional hazards model. Overall survival was calculated from the date of the surgery of primary tumor to death, censoring patients still alive on the last date of follow-up. Glioma-specific survival was calculated from the date of the surgery of primary tumor to glioma caused death, censoring patients still alive or lost from follow-up for other cause on the last date of follow-up. Analyses were conducted with the IBM SPSS 22 software. Results from all other experiments were analyzed using the GraphPad Prism 7 software (La Jolla, USA) using two-tailed, nonparametric Mann-Whitney U test. All the experiments were repeated at least 3 times with triplicates unless stated otherwise.

## Acknowledgments

This work was supported by grants from the Ida Montin, Oskar Öfflund and K. Albin Johansson foundations as well as from the Finnish Cancer Organizations, Jane & Aatos Erkko Foundation and Sigrid Juselius Foundation. M.H. has been supported by the Doctoral Programme in Biomedicine. The Biomedicum Imaging Unit is acknowledged for their assistance in microscopy imaging, Genome Biology Unit (Faculty of Medicine, University of Helsinki, Biocenter Finland) for the 3DHISTECH Pannoramic P250 FLASH II digital slide scanner service. Drs. Outi Rautsi and Kirsi Vuorinen are acknowledged for isolating the patient-derived tumoroids.

## Authors’ Contributions

- **Conception and design:** V. Le Joncour, M. Hyvönen, K. Lehti, P. Laakkonen
- **Development of methodology:** V. Le Joncour, M. Hyvönen, P. Filppu, PS. Turunen, I, Burghardt,
- **Acquisition of data (provided animals, acquired and managed patients, provided facilities, etc.):** V. Le Joncour, M. Hyvönen, P. Filppu, PS. Turunen, O. Tynninen, JE. Jääskeläinen
- **Analysis and interpretation of data (e.g., statistical analysis, biostatistics, computational analysis):** H. Sihto, H. Joensuu, V. Le Joncour, M. Hyvönen, P. Filppu, PS. Turunen
- **Writing, review, and/or revision of the manuscript:** V. Le Joncour, M. Hyvönen, P. Filppu, PS. Turunen, H. Joensuu, O. Tynninen, M. Weller, K. Lehti, P. Laakkonen
- **Administrative, technical, or material support (i.e., reporting or organizing data, constructing databases):** H. Sihto, H. Joensuu, O. Tynninen
- **Study supervision:** V. Le Joncour, I, Burghardt, M. Weller, K. Lehti, P. Laakkonen

## Conflict of Interest

The authors declare that they have no conflict of interest.

